# Spoken language comprehension activates the primary visual cortex

**DOI:** 10.1101/2020.12.02.408765

**Authors:** Anna Seydell-Greenwald, Xiaoying Wang, Elissa Newport, Yanchao Bi, Ella Striem-Amit

## Abstract

Primary visual cortex (V1) is generally thought of as a low-level sensory area that primarily processes basic visual features. However, in congenitally blind individuals, V1 is involved in language processing, with no evidence of major changes in anatomical connectivity that could explain this seemingly drastic functional change. This is at odds with current accounts of neural plasticity, which emphasize the role of connectivity and conserved function in determining a neural tissue’s role even after atypical early experiences. To reconcile what appears to be unprecedented functional reorganization with known accounts of plasticity limitations, we tested whether V1 also responds to spoken language in sighted individuals. Using fMRI, we found that V1 in normally sighted individuals was indeed activated by comprehensible speech as compared to a reversed speech control condition, in a left-lateralized and focal manner. Activation in V1 for language was also significant and comparable for abstract and concrete words, suggesting it was not driven by visual imagery. Last, this activation did not stem from increased attention to the auditory onset of words, excluding general attention accounts. Together these findings suggest that V1 responds to verbal information even in sighted individuals, potentially to predict visual input. This capability might be the basis for the strong V1 language activation observed in people born blind, re-affirming the notion that plasticity is guided by pre-existing connectivity and abilities in the typically developed brain.

**Significance statement:** How flexible is the human brain? Studies of congenitally blind individuals showed that language activates the primary visual cortex. This has been interpreted as evidence for unprecedented functional plasticity from a low-level visual to a language area. To reconcile these findings with known limitations of plasticity based on intrinsic physiology and connectivity, we tested if similar activation can be found in sighted participants. We show that left-lateralized primary visual cortex is activated by spoken language comprehension in sighted individuals, . This suggests that plasticity even in complete blindness from birth is not limitless and is guided by pre-existing connectivity and abilities in the typically-developed brain.

## Introduction

For two decades it has been known that the visual cortex of people born blind can be activated by non-visual inputs, including sound and touch (Sadato et al., 1996; Ptito et al., 2012; Heimler et al., 2014; Renier et al., 2014; Heimler et al., 2015; Cecchetti et al., 2016; Bedny, 2017; Fine and Park, 2018; Bridge and Watkins, 2019). Most functional neuroimaging studies investigating sensory plasticity in the blind have shown that visual association cortex regions perform the same operations (e.g., recognition of objects, script, or motion direction) on input from other sensory modalities (e.g., touch, audition) as they otherwise would on visual input (Renier et al., 2014; Ricciardi et al., 2014; Heimler et al., 2015; Bi et al., 2016; Cecchetti et al., 2016; Peelen and Downing, 2017). These non-visual capacities of the blind are thought to be supported by the typical multisensory connectivity found also in sighted people (Heimler et al., 2015).

In an apparent violation of this rule of conserved function, however, the primary visual cortex (V1) of people born blind has also been shown to be involved in language comprehension and production (Burton et al., 2002a; Röder et al., 2002; Burton et al., 2003; Bedny et al., 2011b; Dietrich et al., 2013; Lane et al., 2015; Abboud et al., 2019). This appears to be a marked deviation from its function in sighted individuals, where V1 is involved in the processing of basic visual features (Hubel and Wiesel, 1962, 1968; Grill-Spector and Malach, 2004; Wandell et al., 2007a). Although evidence for V1 language activation in congenitally blind people is compelling, persuasive evidence for a mechanism by which such extreme functional change from low-level visual to language processing might occur has not been provided to date. Beyond deterioration of the visual pathways (Noppeney et al., 2005; Shimony et al., 2006; Yu et al., 2007; Shu et al., 2009), no drastic differences in anatomical connectivity of the visual cortex have been found between congenitally blind people and sighted controls. Importantly, most recent research suggests that brain organization is strongly determined by anatomical connectivity present already at birth (Mahon and Caramazza, 2011; Hannagan et al., 2015; Heimler et al., 2015; Saygin et al., 2016). This view implies that functional reorganization needs to build on, and is limited by, pre-existing capacities and connections of the available tissue, even in cases of sensory deprivation since birth (Striem-Amit et al., 2018a).

How can findings of language processing in primary visual cortex in the congenitally blind be reconciled with such a view? If the hypothesis of pre-existing anatomical connectivity and its constraints is correct, then to accord with language processing in primary visual cortex in the blind, there must also be language processing in primary visual cortex in sighted people. Evidence pointing in this direction can be found in the observation of V1 activation in several neuroimaging studies of language (e.g., (Bookheimer et al., 1998; Gaillard et al., 2003; Wolmetz et al., 2010; Kovelman et al., 2012). However, these activations could potentially be driven by visual stimulation or imagery evoked by the language stimuli, and they are not discussed in depth in these studies.

To test directly whether language processing engages the primary visual cortex in sighted people, we investigated neural activation in a robust auditory sentence comprehension task, as compared with a low-level control (backward speech, not comprehensible), in 20 neurologically healthy sighted young adults. We also examined and controlled for visual imagery by testing responses in an independent second cohort to auditorily presented abstract words, which are hard to visualize. If language indeed activates primary visual cortex during sentence processing in a typically-developed cohort, such activation may be the basis for the more extreme, previously unaccounted for, plasticity in blindness. This finding would re-affirm the role of pre-existing connectivity and abilities in the intact brain in underlying brain plasticity. Alternatively, the absence of such activation would emphasize the inordinate nature of neural reorganization in blindness.

## Results

To determine whether spoken sentence comprehension activates the visual cortex in the typically developed brain, we compared BOLD activation during blocks of forward versus reverse speech in 20 neurologically healthy sighted young adults (Experiment 1). Playing spoken sentences in reverse renders them incomprehensible while controlling for low-level auditory stimulation, which makes this contrast widely used for studying sentence comprehension (e.g., (Perani et al., 1996; Ahmad et al., 2003; Peña et al., 2003; Gaillard et al., 2007b; Moore-Parks et al., 2010). Participants fixated at screen center throughout the experiment to control for visual stimulation.

In contrasting forward and reverse speech in a whole-brain analysis, a typical left-lateralized fronto-temporal “language” network emerged (**Fig. 1A**), as identified by numerous neuroimaging studies (for reviews, see (Vigneau et al., 2006; Price, 2012)). The primary auditory cortex was not significantly activated because the contrasted conditions are matched in low-level auditory information (see similarly (Seydell-Greenwald et al., 2020)). Importantly, the primary visual cortex was significantly more strongly activated by forward than by reverse speech (**Fig. 1A**). This preference was confirmed by extracting percent signal change from a retinotopically-defined region of interest (ROI) comprising left primary visual cortex (**Fig. 1B**; forward vs. reverse speech, paired t-test, t(19)=4.02, p=0.0004, one-tailed, significant in testing against 0.0125, the Bonferroni-adjusted alpha level for the 4 statistical comparisons performed on the Experiment 1 dataset). The time-course of V1 activation for spoken sentences (**Fig. S1A**) shows a standard stimulus-evoked hemodynamic response.

**Figure 1:**
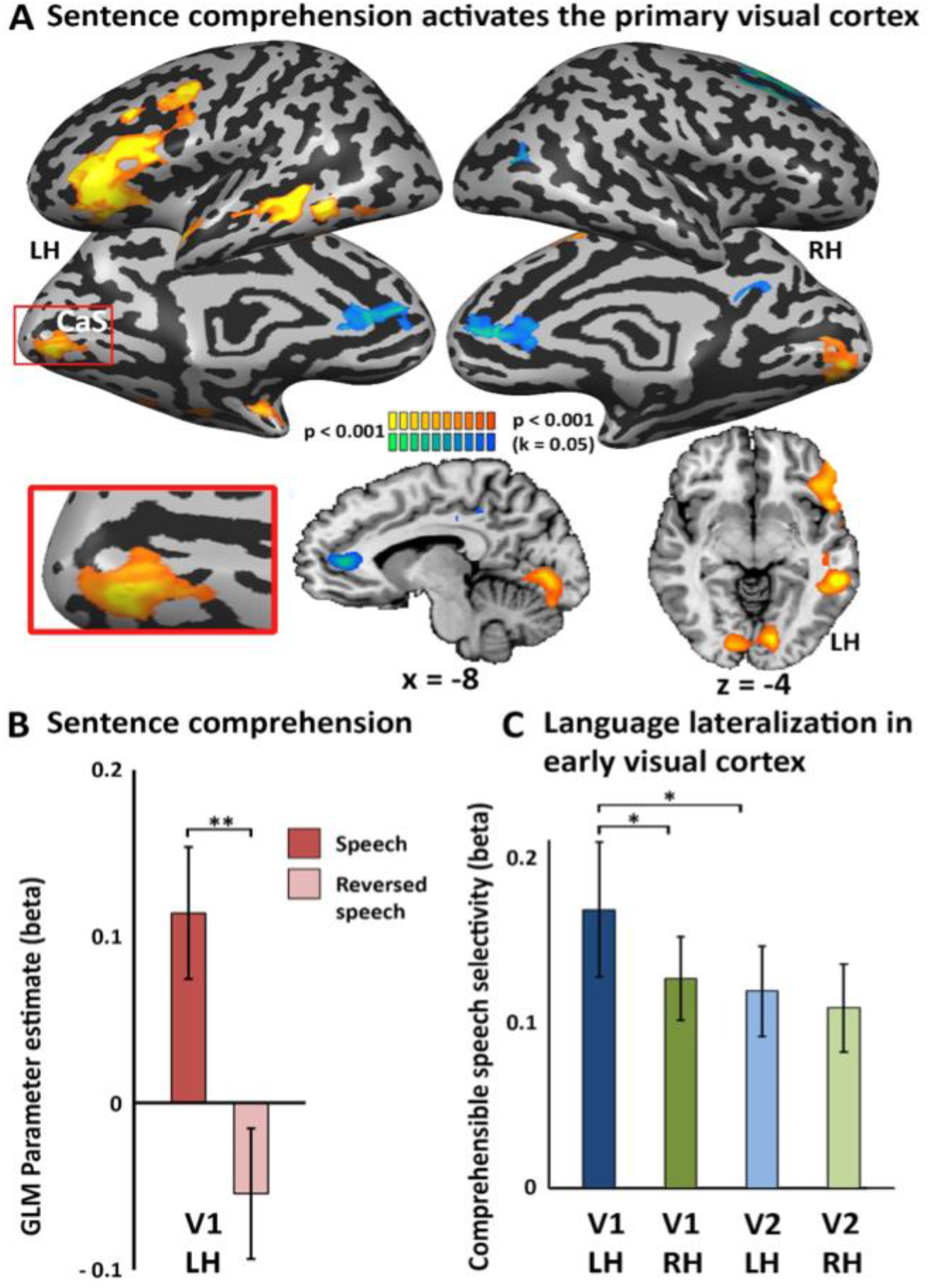
Left primary visual cortex is engaged in spoken language comprehension. **A.** A contrast of comprehensible vs. reversed spoken sentences is shown on brain slices and inflated cortical hemispheres. In addition to the left-lateralized fronto-parieto-temporal language network, significant activation is found in the primary visual cortex. CaS – Calcarine Sulcus. **B.** GLM parameter estimates (betas) were sampled in the left retinotopically-defined primary visual cortex, showing significant selectivity for comprehensible vs. reversed speech. Error bars denote standard error of the mean, **p<0.005 uncorrected, significant with multiple comparisons correction at p<0.05. **C.** Selectivity for comprehensible speech (the beta difference between forward and reversed speech) is higher in the left V1 than in right V1, showing lateralization for language, and compared with V2. Error bars denote standard error of the mean, *p<0.05 uncorrected.

We next compared the forward>reverse speech effect in retinotopically defined primary (V1) and secondary (V2) visual cortex in the left and right hemisphere (**Fig. 1C**) to further investigate the impression from the whole-brain analysis that the activation seemed to be stronger in the left hemisphere and relatively confined to V1. The effect was indeed stronger in left than in right V1, although the difference did not reach statistical significance after multiple comparison correction (left vs. right V1; paired t-test, t(19)=1.98, p=0.031 uncorrected). Moreover, the forward>reverse speech effect was weaker in left V2 than in left V1, although again not statistically significantly after multiple comparison correction (left V1 vs. V2; paired t-test, t(19)=2.06, p=0.027, uncorrected), suggesting that early visual cortex selectivity for language is at least mildly stronger in primary than secondary visual cortex. To test whether the observed V1 activations might reflect increased attention to the onset of auditory stimulation, we repeated the analyses when including a confound condition modelling a short response to the onset of the conditions. This control analysis replicated the main effects (**Fig. S2**), making simple auditory attention effects an unlikely explanation for the V1 language activations.

Could these findings stem from visual imagery, due to the concrete content of the spoken sentences? Although visual imagery is unlikely to explain left-lateralized, focal activation in early visual cortex, we additionally investigated if V1 would show differential activation for abstract (less imaginable) and concrete (easily imaginable) spoken words. Just as for spoken sentence comprehension (Experiment 1 above), whole-brain activation for blocks of abstract words in a separate group of sighted adults (Experiment 2; see also (Striem-Amit et al., 2018b)) included, in addition to vast activation of the temporal lobe and inferior frontal cortex, also significant localized activation in the calcarine sulcus, predominantly in the left hemisphere (**Fig. 2A**). Again, activation was stronger in left than in right V1 (**Fig. 2C**; paired t-test, t(13)=2.77, p=0.016 uncorrected, significant in testing against 0.025, the Bonferroni-adjusted alpha level for the 2 statistical tests performed on the Experiment 2 dataset). Activation time-courses extracted from left V1 resembled the hemodynamic response function for both abstract and concrete words (**Fig. S1C**). Importantly, left V1 activation did not differ between abstract and concrete words (object names; **Fig. 2B**; t(13)=0.48, p=0.65 uncorrected), even though the latter were significantly more imaginable according to behavioral ratings provided by a larger sample of participants (t(9)=1074, p<0.001 uncorrected, significant with correction for multiple comparisons). As in Exp. 1, modelling the potential attention-arousing effect of the auditory onset at the beginning of each block as a nuisance condition did not affect the main findings (**Fig. S3**).

**Figure 2:**
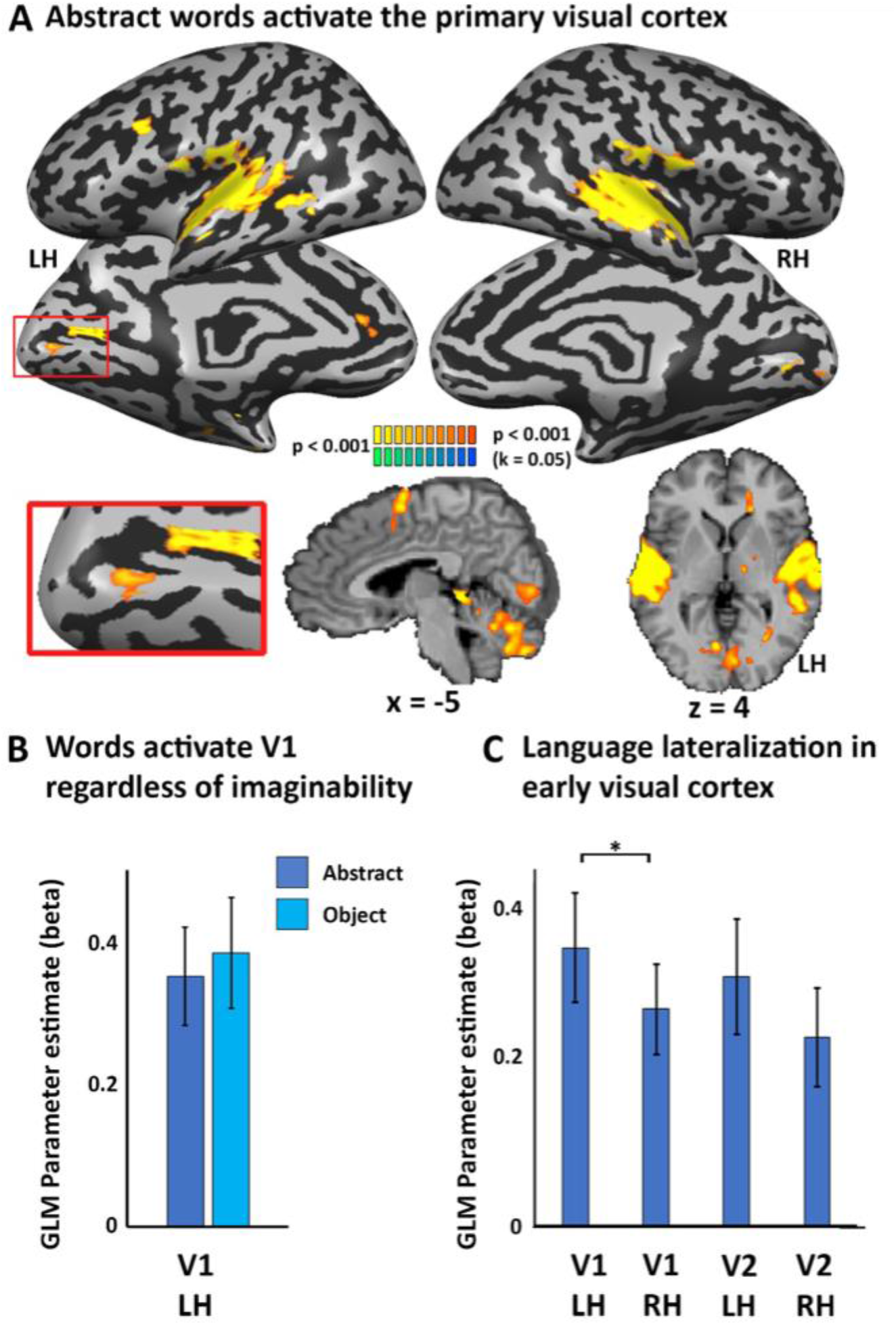
Left primary visual cortex spoken language activation is found for abstract, unimaginable words. **A.** Activation for spoken abstract words is shown on brain slices and inflated cortical hemispheres. In addition to the auditory cortex and inferior frontal cortex, significant activation is found in the primary visual cortex, despite the inability to visually imagine abstract concepts. CaS – Calcarine Sulcus. **B.** GLM parameter estimates (betas) were sampled in the left retinotopically-defined primary visual cortex, showing significant activation for spoken words, which does not differ between abstract and concrete words. **C.** Activation for abstract words is significantly higher in left than right V1, showing lateralization for language. Error bars denote standard error of the mean, *p<0.01 uncorrected, significant with multiple comparisons correction at p<0.05.

Last, an event-related design of spoken words (Experiment 3) allowed us to test whether left V1 activation correlated with imaginability and attentional arousal ratings for spoken words of a variety of imaginable and abstract concept types (Striem-Amit et al., 2018b). No correlation was found between left V1 activation and imaginability ratings (r^2^(54)=0.003, p=0.69 uncorrected) or arousal ratings of these words (r^2^(54)=0.01, p=0.45, uncorrected). Together, these findings suggest that semantic content activation of left V1 does not result from imagery or attention confounds, but rather reflects responses to language comprehension.

## Discussion

The primary visual cortex is widely thought to be a low-level sensory station devoted to the processing of simple visual features (Hubel and Wiesel, 1962, 1968; Grill-Spector and Malach, 2004; Wandell et al., 2007a). However, although still controversial, some recent evidence indicates that it may also receive signals related to higher level non-visual representations, specifically non-visual imagery (Vetter et al., 2014) and working memory (Lawrence et al., 2018). Such atypical activations resemble those observed in the primary visual cortex of people born blind for high-level cognitive functions (Hull and Mason, 1995; Sadato et al., 1996; Sadato et al., 1998; Hamilton et al., 2000; Röder et al., 2000; Burton et al., 2002a; Röder et al., 2002; Amedi et al., 2003; Burton, 2003; Burton et al., 2003; Amedi et al., 2004; Raz et al., 2005; Ofan and Zohary, 2006; Bedny et al., 2011b; Dietrich et al., 2013; Lane et al., 2015; Bedny, 2017; Abboud and Cohen, 2019; Abboud et al., 2019; Vetter et al., 2020) and may provide a possible precursor for such reorganization in the typically developed brain. However, the blind visual cortex has also been implicated in language (Sadato et al., 1996; Burton, 2003; Amedi et al., 2004; Ofan and Zohary, 2006; Bedny, 2017; Abboud and Cohen, 2019; Abboud et al., 2019; Vetter et al., 2020), which has not previously been proposed to meaningfully engage the primary visual cortex in sighted individuals. This apparent lack of a precursor in the sighted has led to the interpretation of V1 language activation in the blind as evidence for extreme functional plasticity.

However, as we show here, this precursor may not be lacking after all. We observed activation for spoken sentences in V1 of typically developed individuals, which showed a preference for comprehensible over incomprehensible speech (**Fig. 1A,B**). Moreover, this activation was left-lateralized (**Fig. 1A,C**), just like the frontotemporal activation typically associated with language tasks, and also like the V1 language activation in blindness (Burton et al., 2002a; Burton, 2003; Ofan and Zohary, 2006; Bedny et al., 2011b). It also seemed to be mostly confined to the primary visual cortex, rather than emerging by feedback cortico-cortical connectivity from visual language areas via higher retinotopic cortical stations (e.g. V2; **Fig. 1C**). The same pattern was evident in response to spoken abstract words in a separate sighted cohort (**Fig. 2**). Together these findings suggest that left-lateralized primary visual cortex responds to semantic information in typically developed sighted adults. These findings have importance for several key issues regarding the multisensory properties of primary visual cortex, the developmental origins of reorganization in the blind brain, and the nature of brain plasticity itself.

Before we discuss these implications, we must first address why our study is the first to highlight V1 language activation in sighted individuals despite the large number of functional neuroimaging studies that have investigated language activation in the typically-developed population. One likely reason is that V1 activation is small relative to that in other regions of the frontotemporal language network, both regarding peak signal change and regarding spatial extent. Thus, depending on the statistical power of the experiment and applied thresholds, V1 activation may not be apparent in all functional neuroimaging studies contrasting speech with silence or non-speech stimuli, although it is apparent in some (Bookheimer et al., 1998; Gaillard et al., 2003; Binder et al., 2004; Zekveld et al., 2006; Holle et al., 2010; Wolmetz et al., 2010; Kovelman et al., 2012; Rämä et al., 2012). An indication that V1 language activation is not uncommon is that meta-analyses on the neurosynth (Yarkoni et al., 2011) and neuroquery (Dockès et al., 2020) platforms reveal converging activation in left early visual cortex for studies on “language” (**Fig. S4**). A look at some of the studies contributing to this converging activation reveals that even if V1 language activation is observed, it is usually either not discussed, explained by imperfectly controlled visual stimulation, or ascribed to top-down attention effects or to visual imagery (O’Leary et al., 1997; Klein et al., 2000; Shaywitz et al., 2001; Cate et al., 2009).

Thus we must next address whether our findings could be explained by confounds such as visual imagery or increased attention to speech. There are several reasons to argue that attention is an unlikely explanation for the V1 activation observed here. First, when attention to sounds activates V1, it serves a role in spatial attention orientation (Cate et al., 2009; Azevedo et al., 2015) and stems from direct connectivity between primary auditory cortices and primarily *peripheral* retinotopic locations of V1 (Rockland and Van Hoesen, 1994; Falchier et al., 2001; Falchier et al., 2002; Rockland and Ojima, 2003; Borra and Rockland, 2011). The activation pattern in comparing forward and reversed speech in our study, in contrast, was not peripherally localized (**Fig. S1B**; ANOVA for an eccentricity effect in left V1 F(2,143)=0.01, p=0.99). Second, no activation was observed in the fronto-parietal attention network (typically bilateral or right-lateralized; (Sturm and Willmes, 2001; Westerhausen et al., 2010; Corbetta and Shulman, 2011; Petersen and Posner, 2012)) in our whole-brain analysis, which would be expected if there were significant attention differences between the conditions (either due to larger top-down attention allocated to processing the comprehensible stimulus, or to larger effort to selectively attend to that stimulus over the scanner noise). Instead, the activation we observed was left-lateralized and, with the exception of the V1 activation, confined to the typical left-lateralized language network, e.g. superior/middle temporal gyrus and inferior frontal gyrus. Moreover, including an additional nuisance regressor to capture attention effects associated with sound onset did not abolish the V1 response in either block-design experiment (**Figs S2 and S3**). Lastly, activation in left V1 was not correlated with the arousal ratings of the heard words in Experiment 3. All this makes it unlikely that attention is the primary driver for the V1 activation we observed.

Similarly, it does not appear to stem from visual imagery. Visual imagery may activate and its content can be decoded from the primary visual cortex (Kosslyn et al., 1995; Chen et al., 1998; Kosslyn et al., 1999; Klein et al., 2000; Slotnick et al., 2005; Cichy et al., 2012; Senden et al., 2019; Ragni et al., 2020). However, imagery responses are stronger in association rather than primary cortex (Reddy et al., 2010; Lee et al., 2012) and are typically bilateral (O’Craven and Kanwisher, 2000; Cichy et al., 2012; Lee et al., 2012). Moreover, V1 involvement has been associated mostly with explicit imagery of high-resolution detail of images (Klein et al., 2000; Kosslyn and Thompson, 2003; Dijkstra et al., 2019). In contrast, our sentence comprehension task did not require explicit imagery or attention to visual detail, and activation was left lateralized and stronger in V1 than V2 (**Fig. 1A,C**). Moreover, we observed the same localized left-lateralized V1 activation in a whole-brain analysis for abstract words (**Fig. 2A,C**), the V1 response did not differ between abstract and concrete words (**Fig. 2B**), and V1 activation was not correlated with imaginability ratings. This pattern of results all but excludes visual imagery as an explanation for the observed V1 activations.

If these V1 language activations are not due merely to attention or imagery, how does linguistic information reach V1 and what role does it play? Primary sensory cortices receive information from multiple cortical and subcortical stations. Specifically, beyond thalamic LGN and pulvinar projections, primary visual cortex receives input from auditory cortices, parietal cortex and other regions including frontal cortex in primates and other mammals (Felleman and Van Essen, 1991; Rockland and Van Hoesen, 1994; Batardiere et al., 1998; Falchier et al., 2001; Falchier et al., 2002; Clavagnier et al., 2004; Hall and Lomber, 2008; Muckli and Petro, 2013; Henschke et al., 2014; Majka et al., 2019; Pennartz et al., 2019). These feedback pathways (Muckli et al., 2015; Pennartz et al., 2019) allow for multisensory integration even in V1 (Ghazanfar and Schroeder, 2006; Murray et al., 2016; Rohe and Noppeney, 2016), along with integration of reward value information (Stănişor et al., 2013; Roth et al., 2020). Theoretically, these cross-modal and higher-level inputs to V1 play a role in predictive coding, whereby predictions of future states and inputs allows for efficient coding and adapting to the everchanging environment (Muckli and Petro, 2013; Pennartz et al., 2019). How language comprehension fits into this framework is uncertain. Language input into V1 may allow integrating contextual information that enables visual cortex to anticipate coming events (Battistoni et al., 2017; Petro et al., 2017; Pennartz et al., 2019). Alternatively, it may play a simpler role in alerting spatial or overall attention, without conveying specific content. It may even be epiphenomenal altogether; our data do not speak directly to these alternative explanations, which will need to be addressed in future work. Importantly, accounts of predictive use of speech information would have to reconcile the level of representation of incoming high-level inputs with the spatial and low-level nature of V1, such that these types of information can be integrated in a meaningful way. Regarding the pathways allowing language to reach V1, the relative confinement of the response we observed to V1 suggests that the typical visual hierarchy (V1->V2-> inferior temporal cortex or vice versa etc.) may not play a role in generating these responses. Similarly, we observed no visual thalamic activation for language selectivity, suggesting that other cortico-cortical pathways may underlie these effects.

Although their role is unclear, the activation patterns observed here suggest an involvement of primary visual cortex in language in a left-lateralized manner in sighted individuals and resemble the findings in people born blind. In people born blind, association visual cortices seem to retain their selectivity towards the same types of information typically processed through vision (e.g. script in the visual word-form area (Reich et al., 2011; Striem-Amit et al., 2012), motion processing in hMT+ (Poirier et al., 2006; Saenz et al., 2008; Ptito et al., 2009; Jiang et al., 2016), and complex visual categories in the ventral visual cortex (Pietrini et al., 2004; Amedi et al., 2007; He et al., 2013; Peelen et al., 2013; Striem-Amit and Amedi, 2014; Peelen and Downing, 2017; van den Hurk et al., 2017; Bola et al., 2020; Mattioni et al., 2020; Ratan Murty et al., 2020). These functions are thought to arise based on existing multisensory processes in and input to these areas, allowing for minimal changes in functional properties (Crollen et al., 2019) even with a change to the dominant input modality (Kupers and Ptito, 2013; Ricciardi et al., 2013; Heimler et al., 2015; Bi et al., 2016). This retention of function in higher visual areas of people born blind contrasts sharply with the apparent dramatic change in the function of their earlier retinotopic visual cortices, including V1, which are activated by high-level cognitive tasks such as language, verbal memory and executive function (Hull and Mason, 1995; Sadato et al., 1998; Hamilton et al., 2000; Röder et al., 2000; Burton et al., 2002a; Röder et al., 2002; Amedi et al., 2003; Burton et al., 2003; Ofan and Zohary, 2006; Bedny et al., 2011b; Dietrich et al., 2013; Lane et al., 2015; Abboud and Cohen, 2019) that do not resemble the original low-level functions of these regions. The fact that stimulating primary visual cortex affects Braille reading (Cohen et al., 1997) and verb generation in the blind (Amedi et al., 2004) suggests that this activation may indeed contribute to language processing.

How does early visual cortex come to process language in the early blind? Even though years have elapsed since the discovery of these atypical activations, with additional evidence suggesting increased functional connectivity between early visual cortex and the inferior frontal lobe in the blind (Liu et al., 2007; Yu et al., 2008; Wang et al., 2013; Burton et al., 2014; Qin et al., 2014; Striem-Amit et al., 2015), a clear mechanism for such remarkable reorganization has not been uncovered. Broadly, with the exception of language-related activation in the congenitally blind visual cortex, cortical plasticity in humans seems to be constrained to relatively small-scale and topographically limited changes in function, even in extreme cases of sensory or motor deprivation from birth (e.g., (Striem-Amit et al., 2018a)). Differences in anatomical connectivity of the visual cortex between congenitally blind people and sighted controls appear to be limited in scope, mostly to the deterioration of the visual pathways in the blind (Noppeney et al., 2005; Shimony et al., 2006; Yu et al., 2007; Shu et al., 2009). Evidently, even in congenital blindness, functional and anatomical connectivity develop largely typically, allowing topographic connectivity for retinotopic areas (Bock et al., 2013; Bock et al., 2015; Striem-Amit et al., 2015) and domain- or category-based connectivity for association cortices (e.g., reviewed in (Heimler et al., 2015). This typical connectivity stems from the fact that large-scale anatomical connectivity, including that of the visual cortex, is largely formed at birth (Burkhalter et al., 1993; Barone et al., 1996; Coogan and Van Essen, 1996; Horton and Hocking, 1996; Takahashi et al., 2012; Arcaro and Livingstone, 2017). Connectivity is principally driven by evolutionarily-constrained genetic cascades that direct brain development (Gomez et al., 2018; Krubitzer and Prescott, 2018) even prior to the onset of visual experience. Therefore, the effects of visual deprivation on anatomical connectivity are limited, and may be restricted to changes in connection weights or synaptic pruning (Innocenti and Price, 2005). Even mechanisms such as decreased pruning of otherwise transient projections (Dehay et al., 1984; Innocenti and Clarke, 1984; Innocenti et al., 1988; Kennedy et al., 1989; Rockland and Van Hoesen, 1994) underlying anatomical rewiring and increased cross-modal connectivity in animal models of blindness (Nicolelis et al., 1991; Karlen et al., 2006; Henschke et al., 2017; Magrou et al., 2017) have not been reported in humans.

An alternative way in which reorganization could occur is the unmasking of existing connections that are not dominant in the presence of sight (Pascual-Leone and Hamilton, 2001). While this proposal does not require massive change in anatomical structure, it assumes that such functions, or at least their predecessor connectivity, exist in the sighted brain. Our findings support this account by demonstrating that primary visual cortex receives language-related information even in sighted people. In early blindness, this normally non-dominant input may become dominant, potentially allowing V1 to functionally contribute to language processing to a much larger extent than it does when its dominant input is visual. Importantly, this explanation of V1 language activation in the early blind does not require pluripotency of visual cortex in the sense of massive changes in function that allow it to perform higher-cognitive functions instead of low-level visual processing (Bedny, 2017). Rather, our data suggest a more conservative explanation of V1’s language recruitment in blindness: little reorganization of V1 structure or function is required to support language recruitment of deprived cortex because it also recruits non-deprived cortex, albeit to a lower extent.

Notably, our findings do not preclude additional reorganization in early onset blindness beyond unmasking. There is abundant evidence for increased activation of visual cortex for language tasks in early onset blindness as compared to sighted people (Burton et al., 2002a; Röder et al., 2002; Burton, 2003; Burton et al., 2003; Bedny et al., 2011b; Lane et al., 2015; Abboud and Cohen, 2019), suggesting a role for local reweighting and better utilization of inputs. Similarly, evidence for differences in the extent of recruitment depending on the timing of blindness onset also suggests that early-onset blindness involves more than unmasking, and that such additional processes are time-sensitive (Cohen et al., 1999; Burton et al., 2002b; Burton et al., 2002a; Burton et al., 2003; Burton and McLaren, 2005; Bedny et al., 2011a; Kanjlia et al., 2018). However, our results reveal a key piece of this puzzle by explaining how language information arrives in early visual cortex. The presence of language responses in the visual cortex of sighted people greatly decreases the requirement for large-scale structural or functional changes to account for this tissue’s recruitment for language in blindness. Further, our evidence brings the inordinate plasticity for language in line with current views of connectivity-driven functional brain organization (Mahon and Caramazza, 2011; Saygin et al., 2011; Hannagan et al., 2015; Heimler et al., 2015; Saygin et al., 2016). Thus we contribute to a unifying explanatory framework for findings in the primary and association cortices in the blind, based on extant non-visual functions of the visual cortex.

In summary, our findings show that the primary visual cortex is engaged in language processing in typically sighted individuals in a localized and left-lateralized manner. Importantly, these findings provide evidence that language-driven visual cortex activation in the blind can be explained without proposing drastic changes to cortical tissue connectivity or function. This suggests that human cortical plasticity is still limited by innate anatomical structures and functional characteristics and is not unconstrained even following extreme changes in early experience.

## Materials and methods

### Participants

#### Experiment 1

Participants were 20 young adults (5 men, ages 18 to 38, mean 21.8 years) with normal or corrected-to-normal vision and no history of neurological disorder from the Georgetown University community. All were native speakers of English and had not been fluent in any other language by the age of 12. All experimental protocols were approved by the institutional review board of Georgetown University Medical Center, in accordance with the Declaration of Helsinki. Participants provided informed consent and were compensated for their time.

#### Experiments 2, 3

Participants were 14 adults with normal or corrected-to-normal vision and no history of neurological disorder (8 men, ages 23 to 66, mean 43.85 years). All were native speakers of Mandarin Chinese. All experimental protocols were approved by the institutional review board of the Department of Psychology at Peking University, China, as well as by the institutional review board of Harvard University, in accordance with the Declaration of Helsinki. Participants provided informed consent and were compensated for their time.

### Experimental Design

#### Experiment 1

The fMRI language task used here was a modified version of an Auditory Description Decision Task used to determine language dominance prior to epilepsy surgery (Gaillard et al., 2007a; Berl et al., 2014). In the Forward Speech condition, participants heard short English sentences (e.g., “A big gray animal is an elephant”) and pushed a button if they considered it a true statement. In the Reverse Speech condition, they heard the same sentences played in reverse (thus rendered incomprehensible) and pushed a button when they heard a soft beep inserted at the end of the utterance. The proportion of correct statements and reverse speech utterances with beeps was 50%. The task was designed to be easy; performance was nearly perfect (median performance at 100% for both tasks, mean 97.2±4.5% for sentence comprehension, mean 99.5±1.1% for beep detection). Each participant completed 2 fMRI runs of 5 min and 48 s duration, each containing 4 30-second blocks of each of 2 experimental conditions (Forward and Reversed Speech, 6 utterances per block) in counterbalanced order, with 12-s silent rest periods at the beginning and end of the run, as well as in between each of the 8 active blocks. Aside from a fixation cross that participants were asked to rest their eyes on throughout the scan, no visual stimulation was provided. The Forward>Reverse activation differences evoked by this task are highly robust and reproducible, making them suitable for localizing language-associated brain areas across development (Olulade et al., 2020) and even in cases of atypical functional organization, such as participants with a history of chronic epilepsy (Berl et al., 2014) or perinatal stroke (Newport et al., 2017). Imaging data were acquired on Georgetown’s research-dedicated 3T Siemens Trio Tim scanner with a 12-channel birdcage head coil. Auditory stimuli were delivered via insert earphones (Sensimetrics S14) worn under ear defenders (Bilsom Thunder T1). Stimulus presentation and response collection (via a Cedrus fiber optics button box) were coordinated by E-Prime 2.0 software. Each of the 2 functional runs contained 100 functional (T2*-weighted) volumes covering the whole brain in 50 slices acquired in descending order and oriented parallel to the AC-PC plane (EPI parameters: TE = 30 ms, TR = 3000 ms, flip angle = 90°, matrix 64×64, slice thickness = mm, distance factor = 7%, resulting in an effective voxel size of 3×3×3 mm^3^). A high-resolution anatomical (T1-weighted) scan was acquired for co-registration (MPRAGE parameters: whole-brain coverage in 176 sagittal slices, TE = 3.5 ms, TR = 2530 ms, TI = 1100 ms, flip angle = 7°, matrix 256×256, voxel size = 1×1×1 mm^3^).

#### Experiments 2, 3

The experiment included presentation of spoken words, each a two-character word in Mandarin Chinese, belonging to eight concept categories: abstract concepts (e.g. “freedom”), concrete everyday object names (e.g. “cup”), and six additional content categories which were not analyzed in the current manuscript (astral/weather phenomena – e.g. “rainbow”, “rain”; scenes – “island”, “beach”; and object features – colors and shapes, e.g. “red”, “square”; see full detail in (Striem-Amit et al., 2018b)). Each category included 10 words whose imaginability and attentional arousal (as well as other measures not used here) were rated on a 7-point scale (Barca et al., 2002) by an independent sample of 45 sighted Chinese participants with similar levels of education. Concrete objects and abstract concepts differed significantly in imaginability (Welch t-test contrast, p< 0.001, but not in arousal p=0.06, uncorrected, after correction for multiple comparisons the difference in imaginability is significant; see full detail in (Striem-Amit et al., 2018b)). During Experiment 2, the participants kept their eyes closed and heard short lists of words in a block design paradigm (8 second blocks with 8 words each, baseline between blocks 8 seconds). Each run began with a 12 sec rest period. Each block contained words from one of the eight concept categories. Experiment 3 was an item-level slow event-related design and was conducted at a different scanning session on the same participants. The stimuli were eight of the ten words of each category from Experiment 2, except for the concrete object names. During each of eight slow event-related runs, the participants heard each word once, in a random order, followed by a 5 second baseline period. In both Experiments 2 and 3 the participant’s task was to detect and respond to semantic catch trials, a fruit name appearing within blocks or as individual events; these blocks/events were removed from further analysis, and runs with more than one missed catch event were excluded. Imaging data were acquired using a Siemens Prisma 3-T scanner with a 20-channel phase-array head coil at the Imaging Center for MRI Research, Peking University. Functional imaging data for Experiment 2 were comprised of four functional runs, each containing 251 continuous whole-brain functional volumes. Functional imaging data for the single-item-level event-related Experiment 3 were comprised of eight functional runs, each containing 209 continuous whole-brain functional volumes. Data was acquired with a simultaneous multi-slice (SMS) sequence supplied by Siemens: slice planes scanned along the rectal gyrus, 64 slices, phase encoding direction from posterior to anterior; 2 mm thickness; 0.2mm gap; multi-band factor = 2; TR = 2000 ms; TE = 30 ms; FA = 90°; matrix size = 112 × 112; FOV = 224 × 224 mm; voxel size = 2 × 2 × 2 mm. T1-weighted anatomical images were acquired for coregistration using a 3D MPRAGE sequence: 192 sagittal slices; 1mm thickness; TR=2530 ms; TE=2.98 ms; inversion time=1100 ms; FA=7°; FOV=256 × 224 mm; voxel size=0.5 × 0.5 × 1 mm, interpolated; matrix size=512 × 448.

### Data Analysis

#### Preprocessing

Imaging data were analyzed using BrainVoyager (BVQX 3.6). Anatomical images were corrected for field inhomogeneities and transformed into Talairach space using 9-parameter affine transformation based on manually identified anatomical landmarks. Functional runs underwent slice time correction, removal of linear trends, and 3D motion correction to the first volume of each run using rigid-body transformation. The first two volumes of each run were discarded to allow for magnetization stabilization. Each run was coregistered with the native-space anatomical image of the same participant using 9-parameter gradient-based alignment, and subsequently warped into Talairach space using the same affine transformation used for warping the anatomical data.

#### Whole-brain group-level analysis

To create group-level activation maps (**Fig. 1A; 2A**), we smoothed the Talairach-warped functional data with a 3D Gaussian kernel of 8 mm FWHM and conducted a hierarchical random effects analysis (RFX GLM; (Friston et al., 1999)). Each experimental condition’s predictor was modeled by convolving the boxcar predictor describing the condition’s time-course with a standard hemodynamic response function (two gamma, peak at 5 s, undershoot peak at 15 s). In addition, the model included nuisance predictors to capture participant- and run-specific effects as well as motion-related effects (using the z-transformed motion estimates generated during preprocessing). During modeling, voxel time courses were normalized using percent signal change transformation and corrected for serial autocorrelations (AR2). Activation maps contrasting the beta values for the different conditions via voxel-wise t-tests were thresholded by applying a single-voxel threshold of p < 0.001 (uncorrected) and running BrainVoyager’s Cluster-Level Statistical Threshold Estimator Plugin to determine a cluster-size threshold corresponding to k < 0.05. Analyses for Exp. 1 and 2 were replicated (**Fig. S2**, **Fig. S3** respectively) with a brief (1TR) condition modelling auditory signal onset at the beginning of each block as a separate predictor, to control for any attention effects elicited by the sound onset after periods of rest.

#### Region-of-interest analyses

Regions-of-interest (ROIs) for the primary and secondary visual cortex (V1 and V2, respectively) were defined from an external localizer (Striem-Amit et al., 2015). The external retinotopy localizer was acquired in a separate group of 14 normally sighted participants using a standard phase-encoded retinotopic mapping protocol, with eccentricity and polar mapping of ring and wedge stimuli, respectively, to measure visual retinotopic mapping (Engel et al., 1994; Sereno et al., 1995; Wandell et al., 2007b; Wandell and Winawer, 2011), delivered during two separate experiments. The experimental detail can be found at (Striem-Amit et al., 2015). Angle (polar) mapping was used to define the borders of V1 and V2 in both hemispheres, used as a ROI to sample activation for the language conditions in the early visual cortices (**Fig. 1B,C; 2B,C**). V1 was further divided to three portions largely representing foveal, middle and peripheral visual fields based on the eccentricity mapping (**Fig. S1B**).

Beta values for each condition were sampled in individuals, and comparisons across conditions within the same ROI were computed with a two-tailed paired t-test. Comparisons across areas were computed based on the subtraction of beta values between forward and reversed speech for each individual, and applying a one-tailed paired t-test between regions, under the prediction that language activation would be localized to the left V1, as seen in blindness (Burton et al., 2002a; Burton, 2003; Burton et al., 2003; Bedny et al., 2011b). In addition, we explored correlations between imaginability and arousal behavioral ratings of the words presented in Experiment 3 and language activation in the left V1 ROI, across all 56 words used in the experiment. T-tests ’p-values are corrected for multiple comparisons using a Bonferroni correction per experiment. Specifically, four statistical comparisons were conducted with Experiment 1: V1 forward vs. reverse speech (**Fig. 1B**), left vs. right V1 and Left V1 vs. V2 (**Fig. 1C**), and the V1 eccentricity effect (**Fig. S1B**). Two statistical comparisons were conducted with Experiment 2: V1 abstract vs. concrete words **(Fig. 2B)**, and left vs. right V1 (**Fig. 2C**). Behavioral data correction for multiple comparisons was based on the number of comparisons made in the context of the whole experimental design behavioral testing; see full detail in (Striem-Amit et al., 2018b). The time-course of activation was also sampled from the left V1 ROI (**Fig. S1A,C**): the averaged percent signal change with relation to condition onset and the standard errors were calculated for each condition and plotted for each time point.

## Acknowledgements

This work was supported by a KL2 grant (to A.S.-G.) from NIH TL1TR001431 Georgetown–Howard Universities Clinical and Translational Science Award, the Georgetown/MedStar Center for Brain Plasticity and Recovery support (to A.S-G), the National Natural Science Foundation of China (31500882 to X.Y.W., 31671128 to Y.B.); the Fundamental Research Funds for the Central Universities (2017XTCX04, to Y.B.) and Interdisciplinary Research Funds of Beijing Normal University (to Y.B.), by the European Union’s Horizon 2020 Research and Innovation Programme under Marie Sklodowska-Curie Grant Agreement 654837 (to E.S-A.); and the Edwin H. Richard and Elisabeth Richard von Matsch Distinguished Professorship in Neurological Diseases (to E.S.-A.).

## Supplementary Material

**Figure S1:**
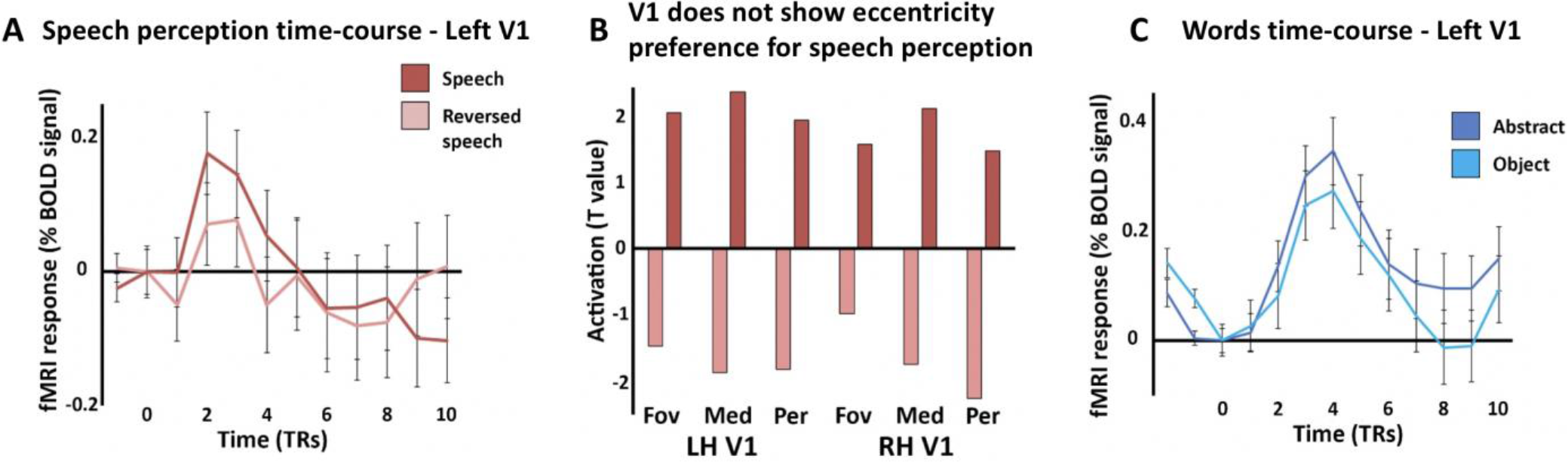
Activation for speech comprehension in primary visual cortex. A. Time course of activation from Experiment 1 was sampled from retinotopic left V1, showing typical BOLD-shaped response in V1 for speech, which is higher for forward as compared to reversed speech. B. GLM parameter estimates (betas) were sampled in the left retinotopically-defined primary visual cortex divided based on eccentricity, showing that the activation for forward speech does not differ between foveal, middle and peripheral-representing V1 sections. C. Time course of activation from Experiment 2 was sampled from retinotopic left V1, showing typical BOLD-shaped response in V1 for abstract and concrete words.

**Figure S2:**
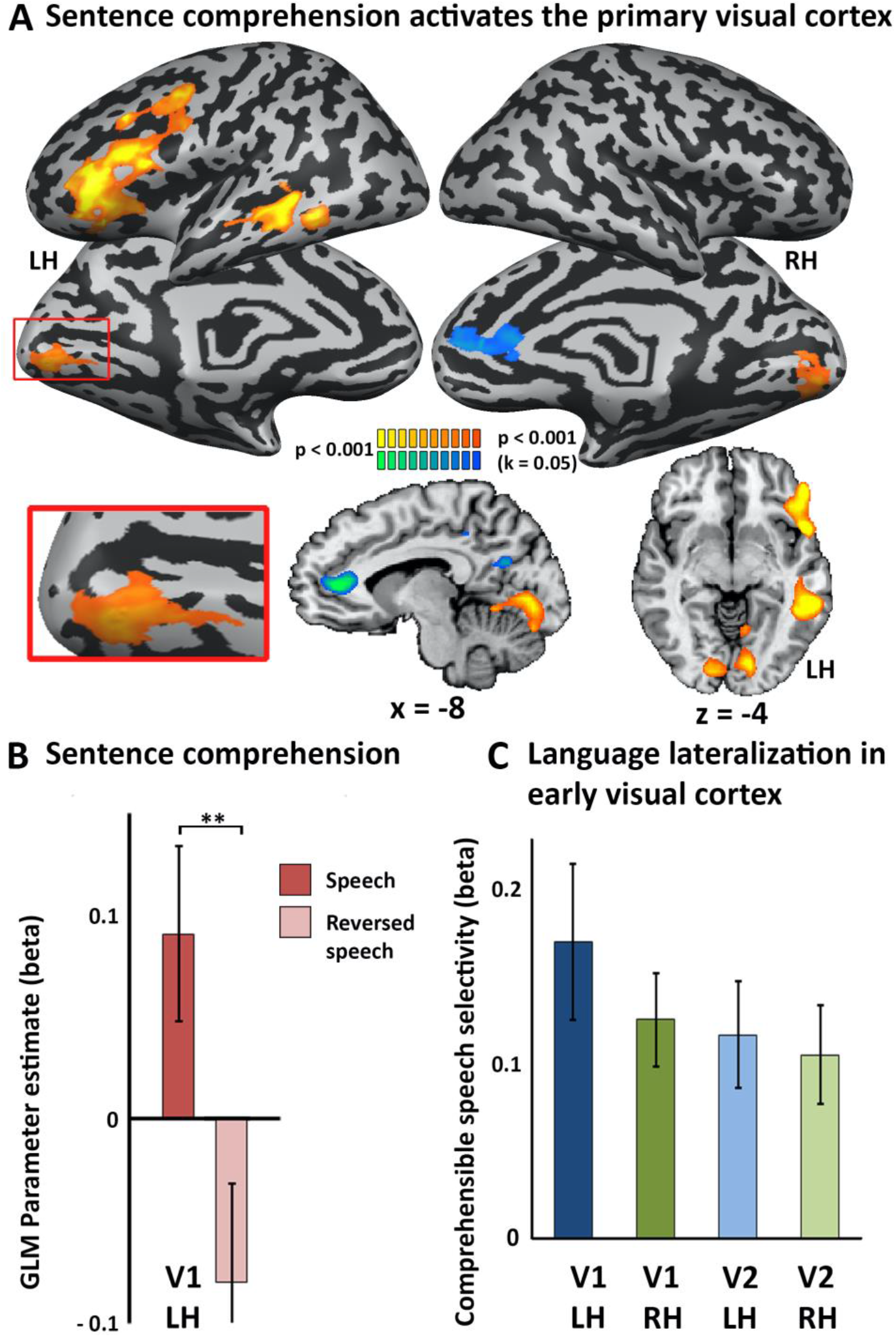
Left primary visual cortex is engaged in spoken language comprehension when including a “block onset” predictor to control for bottom-up attention effects. This figure is provided for comparison with Figure 1. The underlying analyses are the same except for inclusion of an additional nuisance predictor in the GLM to capture the bottom-up attention effects that might occur at the beginning of each block, at the onset of auditory stimulation. Even after including this additional predictor, a left-lateralized fronto-temporal language network is clearly evident (A), left V1 activation is significantly stronger for comprehensible forward than incomprehensible reverse speech (B), and the forward>reverse effect is stronger in left V1 compared to right V1 and left V2 (C). **p<0.005 uncorrected, significant with multiple comparisons correction at p<0.05.

**Figure S3:**
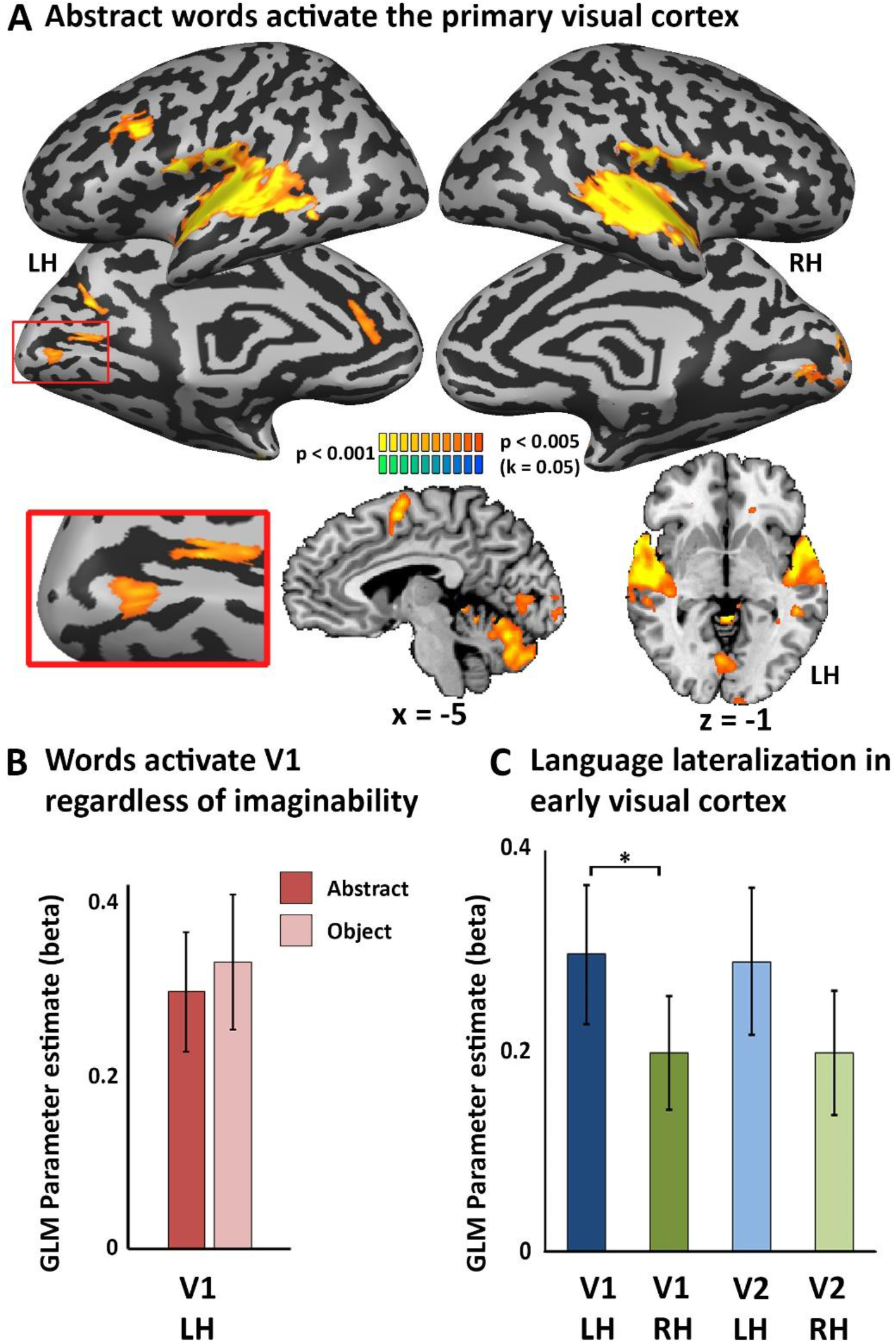
Left primary visual cortex spoken language activation is found for abstract, unimaginable words when including a “block onset” predictor to control for bottom-up attention effects. This figure is provided for comparison with Figure 2. The underlying analyses are the same except for inclusion of an additional nuisance predictor in the GLM to capture the bottom-up attention effects that might occur at the beginning of each block, at the onset of auditory stimulation. Even after including this additional predictor, activation of the primary visual cortex for abstract words is clearly evident (A), with no difference in response between abstract and concrete (object name) words (B; p=0.60), and the response to abstract words is stronger in left V1 compared to right V1 (C). *p<0.01 uncorrected, significant with multiple comparisons correction at p<0.05.

**Figure S2:**
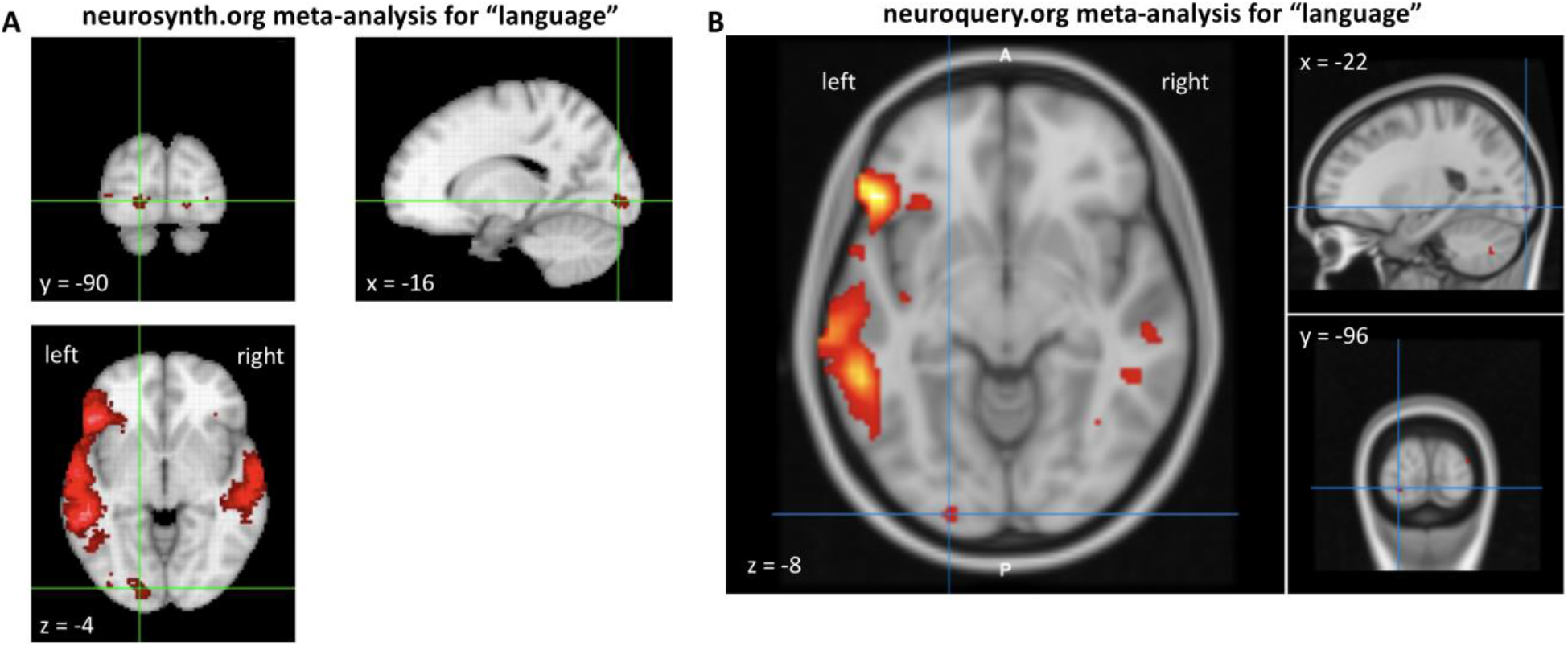
Left-lateralized early visual activation in “language” meta-analyses. A. Screenshot of the results of a coordinate-based meta-analysis using the neurosynth.org platform (Yarkoni et al., 2011) showing a left early visual cluster. The map is based on the activation coordinates reported in 1101 functional neuroimaging studies based on a two-way ANOVA testing whether activation at the voxel occurs more consistently in studies mentioning the term “language” with high frequency in their full text than in studies that do not. The map is thresholded to limit the false discovery rate to 1% (FDR<0.01). The left early visual cortex activation highlighted by the crosshairs is highly significant (z=4.4). B. Screenshot of the predicted activation for “language” using the neuroquery.org platform (Dockès et al., 2020) showing a left early visual cluster. The map is derived using a reduced-rank linear regression model on the activation coordinates reported in a training corpus of 14,000 neuroimaging studies. The “language” map was automatically generated using Neuroquery and included mainly contribution from 73 neuroimaging studies associated with the term “language” and several additional studies on related terms such as “speech” or “comprehension”. The z-value at the voxel highlighted by the crosshairs is z=3.9.

## References

Abboud S, Cohen L (2019) Distinctive Interaction Between Cognitive Networks and the Visual Cortex in Early Blind Individuals. Cereb Cortex.

Abboud S, Engemann D-A, Cohen L (2019) Semantic coding in the occipital cortex of early blind individuals. bioRxiv:539437.

Ahmad Z, Balsamo LM, Sachs BC, Xu B, Gaillard WD (2003) Auditory comprehension of language in young children. Neural networks identified with fMRI 60:1598–1605.

Amedi A, Raz N, Pianka P, Malach R, Zohary E (2003) Early ‘visual’ cortex activation correlates with superior verbal memory performance in the blind. Nat Neurosci 6:758–766.

Amedi A, Floel A, Knecht S, Zohary E, Cohen LG (2004) Transcranial magnetic stimulation of the occipital pole interferes with verbal processing in blind subjects. Nat Neurosci 7:1266–1270.

Amedi A, Stern WM, Camprodon JA, Bermpohl F, Merabet L, Rotman S, Hemond C, Meijer P, Pascual-Leone A (2007) Shape conveyed by visual-to-auditory sensory substitution activates the lateral occipital complex. Nat Neurosci 10:687–689.

Arcaro MJ, Livingstone MS (2017) A hierarchical, retinotopic proto-organization of the primate visual system at birth. eLife 6:e26196.

Azevedo FA, Ortiz-Rios M, Li Q, Logothetis NK, Keliris GA (2015) A Potential Role of Auditory Induced Modulations in Primary Visual Cortex. Multisens Res 28:331–349.

Barca L, Burani C, Arduino LS (2002) Word naming times and psycholinguistic norms for Italian nouns. Behav Res Methods Instrum Comput 34:424–434.

Barone P, Dehay C, Berland M, Kennedy H (1996) Role of directed growth and target selection in the formation of cortical pathways: Prenatal development of the projection of area V2 to area V4 in the monkey. Journal of Comparative Neurology 374:1–20.

Batardiere A, Barone P, Dehay C, Kennedy H (1998) Area-specific laminar distribution of cortical feedback neurons projecting to cat area 17: quantitative analysis in the adult and during ontogeny. J Comp Neurol 396:493–510.

Battistoni E, Stein T, Peelen MV (2017) Preparatory attention in visual cortex. Annals of the New York Academy of Sciences.

Bedny M (2017) Evidence from Blindness for a Cognitively Pluripotent Cortex. Trends in Cognitive Sciences 21:637–648.

Bedny M, Pascual-Leone A, Dravida S, Saxe R (2011a) A sensitive period for language in the visual cortex: Distinct patterns of plasticity in congenitally versus late blind adults. Brain Lang.

Bedny M, Pascual-Leone A, Dodell-Feder D, Fedorenko E, Saxe R (2011b) Language processing in the occipital cortex of congenitally blind adults. Proc Natl Acad Sci U S A 108:4429–4434.

Berl MM, Zimmaro LA, Khan OI, Dustin I, Ritzl E, Duke ES, Sepeta LN, Sato S, Theodore WH, Gaillard WD (2014) Characterization of atypical language activation patterns in focal epilepsy. Ann Neurol 75:33–42.

Bi Y, Wang X, Caramazza A (2016) Object Domain and Modality in the Ventral Visual Pathway. Trends in Cognitive Sciences.

Binder JR, Liebenthal E, Possing ET, Medler DA, Ward BD (2004) Neural correlates of sensory and decision processes in auditory object identification. Nature Neuroscience 7:295–301.

Bock AS, Saenz M, Tungaraza R, Boynton GM, Bridge H, Fine I (2013) Visual callosal topography in the absence of retinal input. Neuroimage.

Bock AS, Binda P, Benson NC, Bridge H, Watkins KE, Fine I (2015) Resting-State Retinotopic Organization in the Absence of Retinal Input and Visual Experience. The Journal of Neuroscience 35:12366–12382.

Bola Ł, Yang H, Caramazza A, Bi Y (2020) Preference for animate domain sounds in the fusiform gyrus of blind individuals is modulated by shape-action mapping. bioRxiv:2020.2006.2020.162917.

Bookheimer SY, Zeffiro TA, Blaxton TA, Gaillard WD, Malow B, Theodore WH (1998) Regional cerebral blood flow during auditory responsive naming: evidence for cross-modality neural activation. Neuroreport 9:2409–2413.

Borra E, Rockland KS (2011) Projections to early visual areas v1 and v2 in the calcarine fissure from parietal association areas in the macaque. Frontiers in neuroanatomy 5:35–35.

Bridge H, Watkins KE (2019) Structural and functional brain reorganisation due to blindness: The special case of bilateral congenital anophthalmia. Neuroscience & Biobehavioral Reviews 107:765–774.

Burkhalter A, Bernardo KL, Charles V (1993) Development of local circuits in human visual cortex. J Neurosci 13:1916–1931.

Burton H (2003) Visual cortex activity in early and late blind people. J Neurosci 23:4005–4011.

Burton H, McLaren DG (2005) Visual cortex activation in late-onset, Braille naive blind individuals: An fMRI study during semantic and phonological tasks with heard words. Neurosci Lett.

Burton H, Diamond JB, McDermott KB (2003) Dissociating cortical regions activated by semantic and phonological tasks: a FMRI study in blind and sighted people. J Neurophysiol 90:1965–1982. Epub 2003 Jun 1964.

Burton H, Snyder AZ, Raichle ME (2014) Resting state functional connectivity in early blind humans. Front Syst Neurosci 8:51.

Burton H, Snyder AZ, Diamond JB, Raichle ME (2002a) Adaptive changes in early and late blind: a FMRI study of verb generation to heard nouns. J Neurophysiol 88:3359–3371.

Burton H, Snyder AZ, Conturo TE, Akbudak E, Ollinger JM, Raichle ME (2002b) Adaptive changes in early and late blind: a fMRI study of Braille reading. J Neurophysiol 87:589–607.

Cate AD, Herron TJ, Yund EW, Stecker GC, Rinne T, Kang X, Petkov CI, Disbrow EA, Woods DL (2009) Auditory attention activates peripheral visual cortex. PLoS ONE 4:e4645.

Cecchetti L, Kupers R, Ptito M, Pietrini P, Ricciardi E (2016) Are Supramodality and Cross-Modal Plasticity the Yin and Yang of Brain Development? From Blindness to Rehabilitation. Frontiers in Systems Neuroscience 10.

Chen W, Kato T, Zhu XH, Ogawa S, Tank DW, Ugurbil K (1998) Human primary visual cortex and lateral geniculate nucleus activation during visual imagery. Neuroreport 9:3669–3674.

Cichy RM, Heinzle J, Haynes JD (2012) Imagery and perception share cortical representations of content and location. Cereb Cortex 22:372–380.

Clavagnier S, Falchier A, Kennedy H (2004) Long-distance feedback projections to area V1: implications for multisensory integration, spatial awareness, and visual consciousness. Cogn Affect Behav Neurosci 4:117–126.

Cohen LG, Weeks RA, Sadato N, Celnik P, Ishii K, Hallett M (1999) Period of susceptibility for cross-modal plasticity in the blind. Ann Neurol 45:451–460.

Cohen LG, Celnik P, Pascual-Leone A, Corwell B, Falz L, Dambrosia J, Honda M, Sadato N, Gerloff C, Catala MD, Hallett M (1997) Functional relevance of cross-modal plasticity in blind humans. Nature 389:180–183.

Coogan TA, Van Essen DC (1996) Development of connections within and between areas V1 and V2 of macaque monkeys. Journal of Comparative Neurology 372:327–342.

Corbetta M, Shulman GL (2011) Spatial neglect and attention networks. Annu Rev Neurosci 34:569–599.

Crollen V, Lazzouni L, Bellemare A, Rezk M, Lepore F, Noel M-P, Seron X, Collignon O (2019) Recruitment of occipital cortex by arithmetic processing follows computational bias in early blind. Neuroimage 186:549–556.

Dehay C, Bullier J, Kennedy H (1984) Transient projections from the fronto-parietal and temporal cortex to areas 17, 18 and 19 in the kitten. Exp Brain Res 57:208–212.

Dietrich S, Hertrich I, Ackermann H (2013) Ultra-fast speech comprehension in blind subjects engages primary visual cortex, fusiform gyrus, and pulvinar -- a functional magnetic resonance imaging (fMRI) study. BMC Neurosci 14:74.

Dijkstra N, Bosch S, Gerven M (2019) Shared Neural Mechanisms of Visual Perception and Imagery. Trends in Cognitive Sciences 23.

Dockès J, Poldrack RA, Primet R, Gözükan H, Yarkoni T, Suchanek F, Thirion B, Varoquaux G (2020) NeuroQuery, comprehensive meta-analysis of human brain mapping. eLife 9:e53385.

Engel SA, Rumelhart DE, Wandell BA, Lee AT, Glover GH, Chichilnisky EJ, Shadlen MN (1994) fMRI of human visual cortex. Nature 369:525.

Falchier A, Renaud L, Barone P, Kennedy H (2001) Extensive projections from the primary auditory cortex and polisensory area STP to peripheral area V1 in the macaque. Society for Neuroscience Abstracts 31:511–521.

Falchier A, Clavagnier S, Barone P, Kennedy H (2002) Anatomical evidence of multimodal integration in primate striate cortex. J Neurosci 22:5749–5759.

Felleman DJ, Van Essen DC (1991) Distributed hierarchical processing in the primate cerebral cortex. Cereb Cortex 1:1–47.

Fine I, Park J-M (2018) Blindness and Human Brain Plasticity. Annual Review of Vision Science 4:337–356.

Friston KJ, Holmes AP, Worsley KJ (1999) How many subjects constitute a study? Neuroimage 10:1–5.

Gaillard WD, Sachs BC, Whitnah JR, Ahmad Z, Balsamo LM, Petrella JR, Braniecki SH, McKinney CM, Hunter K, Xu B, Grandin CB (2003) Developmental aspects of language processing: fMRI of verbal fluency in children and adults. Human Brain Mapping 18:176–185.

Gaillard WD, Berl MM, Moore EN, Ritzl EK, Rosenberger LR, Weinstein SL, Conry JA, Pearl PL, Ritter FF, Sato S, Vezina LG, Vaidya CJ, Wiggs E, Fratalli C, Risse G, Ratner NB, Gioia G, Theodore WH (2007a) Atypical language in lesional and nonlesional complex partial epilepsy. Neurology 69:1761–1771.

Gaillard WD, Berl MM, Moore EN, Ritzl EK, Rosenberger LR, Weinstein SL, Conry JA, Pearl PL, Ritter FF, Sato S, Vezina LG, Vaidya CJ, Wiggs E, Fratalli C, Risse G, Ratner NB, Gioia G, Theodore WH (2007b) Atypical language in lesional and nonlesional complex partial epilepsy. Neurology 69:1761–1771.

Ghazanfar AA, Schroeder CE (2006) Is neocortex essentially multisensory? Trends in Cognitive Sciences 10:278–285.

Gomez J, Zhen Z, Weiner K (2018) Human visual cortex is organized along two genetically opposed hierarchical gradients with unique developmental and evolutionary origins. BioRxiv.

Grill-Spector K, Malach R (2004) The human visual cortex. Annu Rev Neurosci 27:649–677.

Hall AJ, Lomber SG (2008) Auditory cortex projections target the peripheral field representation of primary visual cortex. Exp Brain Res 190:413–430.

Hamilton R, Keenan JP, Catala M, Pascual-Leone A (2000) Alexia for Braille following bilateral occipital stroke in an early blind woman. Neuroreport 11:237–240.

Hannagan T, Amedi A, Cohen L, Dehaene-Lambertz G, Dehaene S (2015) Origins of the specialization for letters and numbers in ventral occipitotemporal cortex. Trends in Cognitive Sciences.

He C, Peelen MV, Han Z, Lin N, Caramazza A, Bi Y (2013) Selectivity for large nonmanipulable objects in scene-selective visual cortex does not require visual experience. Neuroimage 79:1–9.

Heimler B, Weisz N, Collignon O (2014) Revisiting the adaptive and maladaptive effects of crossmodal plasticity. Neuroscience 283:44–63.

Heimler B, Striem-Amit E, Amedi A (2015) Origins of task-specific sensory-independent organization in the visual and auditory brain: neuroscience evidence, open questions and clinical implications. Curr Opin Neurobiol 35:169–177.

Henschke J, Noesselt T, Scheich H, Budinger E (2014) Possible anatomical pathways for short-latency multisensory integration processes in primary sensory cortices. Brain Structure and Function:1–23.

Henschke JU, Oelschlegel AM, Angenstein F, Ohl FW, Goldschmidt J, Kanold PO, Budinger E (2017) Early sensory experience influences the development of multisensory thalamocortical and intracortical connections of primary sensory cortices. Brain Struct Funct.

Holle H, Obleser J, Rueschemeyer S-A, Gunter TC (2010) Integration of iconic gestures and speech in left superior temporal areas boosts speech comprehension under adverse listening conditions. Neuroimage 49:875–884.

Horton JC, Hocking DR (1996) An adult-like pattern of ocular dominance columns in striate cortex of newborn monkeys prior to visual experience. J Neurosci 16:1791–1807.

Hubel DH, Wiesel TN (1962) Receptive fields, binocular interaction and functional architecture in the cat’s visual cortex. The Journal of physiology 160:106–154.

Hubel DH, Wiesel TN (1968) Receptive fields and functional architecture of monkey striate cortex. The Journal of Physiology 195:215–243.

Hull T, Mason H (1995) Performance of blind-children on digit-span tests. Journal of Visual Impairment and Blindness 89:166–169.

Innocenti GM, Clarke S (1984) Bilateral transitory projection to visual areas from auditory cortex in kittens. Brain Res 316:143–148.

Innocenti GM, Price DJ (2005) Exuberance in the development of cortical networks. Nat Rev Neurosci 6:955–965.

Innocenti GM, Berbel P, Clarke S (1988) Development of projections from auditory to visual areas in the cat. J Comp Neurol 272:242–259.

Jiang F, Stecker GC, Boynton GM, Fine I, Van Kemenade B, Collignon O, Burton H (2016) Early Blindness Results in Developmental Plasticity for Auditory Motion Processing within Auditory and Occipital Cortex. Frontiers in Human Neuroscience.

Kanjlia S, Pant R, Bedny M (2018) Sensitive Period for Cognitive Repurposing of Human Visual Cortex. Cerebral Cortex:bhy280–bhy280.

Karlen SJ, Kahn DM, Krubitzer L (2006) Early blindness results in abnormal corticocortical and thalamocortical connections. Neuroscience 142:843–858.

Kennedy H, Bullier J, Dehay C (1989) Transient projection from the superior temporal sulcus to area 17 in the newborn macaque monkey. Proc Natl Acad Sci U S A 86:8093–8097.

Klein I, Paradis AL, Poline JB, Kosslyn SM, Le Bihan D (2000) Transient activity in the human calcarine cortex during visual-mental imagery: an event-related fMRI study. J Cogn Neurosci 12 Suppl 2:15–23.

Kosslyn SM, Thompson WL (2003) When is early visual cortex activated during visual mental imagery? Psychol Bull 129:723–746.

Kosslyn SM, Thompson WL, Kim IJ, Alpert NM (1995) Topographical representations of mental images in primary visual cortex. Nature 378:496–498.

Kosslyn SM, Pascual-Leone A, Felician O, Camposano S, Keenan JP, Thompson WL, Ganis G, Sukel KE, Alpert NM (1999) The role of area 17 in visual imagery: convergent evidence from PET and rTMS. Science 284:167–170.

Kovelman I, Norton ES, Christodoulou JA, Gaab N, Lieberman DA, Triantafyllou C, Wolf M, Whitfield-Gabrieli S, Gabrieli JDE (2012) Brain Basis of Phonological Awareness for Spoken Language in Children and Its Disruption in Dyslexia. Cerebral Cortex 22:754–764.

Krubitzer LA, Prescott TJ (2018) The Combinatorial Creature: Cortical Phenotypes within and across Lifetimes. Trends in Neurosciences 41:744–762.

Kupers R, Ptito M (2013) Compensatory plasticity and cross-modal reorganization following early visual deprivation. Neurosci Biobehav Rev.

Lane C, Kanjlia S, Omaki A, Bedny M (2015) “Visual” Cortex of Congenitally Blind Adults Responds to Syntactic Movement. The Journal of Neuroscience 35:12859–12868.

Lawrence SJD, van Mourik T, Kok P, Koopmans PJ, Norris DG, de Lange FP (2018) Laminar Organization of Working Memory Signals in Human Visual Cortex. Current Biology 28:3435–3440.e3434.

Lee SH, Kravitz DJ, Baker CI (2012) Disentangling visual imagery and perception of real-world objects. Neuroimage 59:4064–4073.

Liu Y, Yu C, Liang M, Li J, Tian L, Zhou Y, Qin W, Li K, Jiang T (2007) Whole brain functional connectivity in the early blind. Brain 130:2085–2096.

Magrou L, Barone P, Markov NT, Killackey H, Giroud P, Berland M, Knoblauch K, Dehay C, Kennedy H (2017) Cortical Connectivity In A Macaque Model Of Congenital Blindness. bioRxiv.

Mahon BZ, Caramazza A (2011) What drives the organization of object knowledge in the brain? Trends Cogn Sci 15:97–103.

Majka P, Rosa MGP, Bai S, Chan JM, Huo B-X, Jermakow N, Lin MK, Takahashi YS, Wolkowicz IH, Worthy KH, Rajan R, Reser DH, Wójcik DK, Okano H, Mitra PP (2019) Unidirectional monosynaptic connections from auditory areas to the primary visual cortex in the marmoset monkey. Brain structure & function 224:111–131.

Mattioni S, Rezk M, Battal C, Bottini R, Cuculiza Mendoza KE, Oosterhof NN, Collignon O (2020) Categorical representation from sound and sight in the ventral occipito-temporal cortex of sighted and blind. eLife 9:e50732.

Moore-Parks EN, Burns EL, Bazzill R, Levy S, Posada V, Müller RA (2010) An fMRI study of sentence-embedded lexical-semantic decision in children and adults. Brain Lang 114:90–100.

Muckli L, Petro LS (2013) Network interactions: non-geniculate input to V1. Current Opinion in Neurobiology 23:195–201.

Muckli L, De Martino F, Vizioli L, Petro Lucy S, Smith Fraser W, Ugurbil K, Goebel R, Yacoub E (2015) Contextual Feedback to Superficial Layers of V1. Current Biology 25:2690–2695.

Murray MM, Thelen A, Thut G, Romei V, Martuzzi R, Matusz PJ (2016) The multisensory function of the human primary visual cortex. Neuropsychologia 83:161–169.

Newport EL, Landau B, Seydell-Greenwald A, Turkeltaub PE, Chambers CE, Dromerick AW, Carpenter J, Berl MM, Gaillard WD (2017) Revisiting Lenneberg’s Hypotheses About Early Developmental Plasticity: Language Organization After Left-Hemisphere Perinatal Stroke. Biolinguistics (Nicos) 11:407–422.

Nicolelis MA, Chapin JK, Lin RC (1991) Neonatal whisker removal in rats stabilizes a transient projection from the auditory thalamus to the primary somatosensory cortex. Brain Res 567:133–139.

Noppeney U, Friston KJ, Ashburner J, Frackowiak R, Price CJ (2005) Early visual deprivation induces structural plasticity in gray and white matter. Curr Biol 15:R488–490.

O’Craven KM, Kanwisher N (2000) Mental imagery of faces and places activates corresponding stiimulus-specific brain regions. J Cogn Neurosci 12:1013–1023.

O’Leary DS, Andreasen NC, Hurtig RR, Torres IJ, Flashman LA, Kesler ML, Arndt SV, Cizadlo TJ, Ponto LLB, Watkins GL, Hichwa RD (1997) Auditory and visual attention assessed with PET. Human Brain Mapping 5:422–436.

Ofan RH, Zohary E (2006) Visual Cortex Activation in Bilingual Blind Individuals during Use of Native and Second Language. Cereb Cortex 17:1249–1259.

Olulade OA, Seydell-Greenwald A, Chambers CE, Turkeltaub PE, Dromerick AW, Berl MM, Gaillard WD, Newport EL (2020) The neural basis of language development: Changes in lateralization over age. Proceedings of the National Academy of Sciences:201905590.

Pascual-Leone A, Hamilton R (2001) The metamodal organization of the brain. Prog Brain Res 134:427–445.

Peelen MV, Downing PE (2017) Category selectivity in human visual cortex: Beyond visual object recognition. Neuropsychologia.

Peelen MV, Bracci S, Lu X, He C, Caramazza A, Bi Y (2013) Tool Selectivity in Left Occipitotemporal Cortex Develops without Vision. Journal of Cognitive Neuroscience:1–10.

Peña M, Maki A, Kovacić D, Dehaene-Lambertz G, Koizumi H, Bouquet F, Mehler J (2003) Sounds and silence: an optical topography study of language recognition at birth. Proc Natl Acad Sci U S A 100:11702–11705.

Pennartz CMA, Dora S, Muckli L, Lorteije JAM (2019) Towards a Unified View on Pathways and Functions of Neural Recurrent Processing. Trends in Neurosciences 42:589–603.

Perani D, Dehaene S, Grassi F, Cohen L, Cappa SF, Dupoux E, Fazio F, Mehler J (1996) Brain processing of native and foreign languages. Neuroreport 7:2439–2444.

Petersen SE, Posner MI (2012) The attention system of the human brain: 20 years after. Annu Rev Neurosci 35:73–89.

Petro LS, Paton AT, Muckli L (2017) Contextual modulation of primary visual cortex by auditory signals. Philosophical Transactions of the Royal Society B: Biological Sciences 372.

Pietrini P, Furey ML, Ricciardi E, Gobbini MI, Wu WH, Cohen L, Guazzelli M, Haxby JV (2004) Beyond sensory images: Object-based representation in the human ventral pathway. Proc Natl Acad Sci U S A 101:5658–5663. Epub 2004 Apr 5652.

Poirier CC, Collignon O, Scheiber C, Renier L, Vanlierde A, Tranduy D, Veraart C, De Volder AG (2006) Auditory motion perception activates visual motion areas in early blind subjects. Neuroimage 31:279–285.

Price CJ (2012) A review and synthesis of the first 20 years of PET and fMRI studies of heard speech, spoken language and reading. Neuroimage.

Ptito M, Matteau I, Gjedde A, Kupers R (2009) Recruitment of the middle temporal area by tactile motion in congenital blindness. Neuroreport 20:543–547.

Ptito M, Matteau I, Zhi Wang A, Paulson OB, Siebner HR, Kupers R (2012) Crossmodal Recruitment of the Ventral Visual Stream in Congenital Blindness. Neural Plasticity 2012.

Qin W, Xuan Y, Liu Y, Jiang T, Yu C (2014) Functional Connectivity Density in Congenitally and Late Blind Subjects. Cerebral Cortex.

Ragni F, Tucciarelli R, Andersson P, Lingnau A (2020) Decoding stimulus identity in occipital, parietal and inferotemporal cortices during visual mental imagery. Cortex.

Rämä P, Relander-Syrjänen K, Carlson S, Salonen O, Kujala T (2012) Attention and semantic processing during speech: an fMRI study. Brain Lang 122:114–119.

Ratan Murty NA, Teng S, Beeler D, Mynick A, Oliva A, Kanwisher N (2020) Visual experience is not necessary for the development of face-selectivity in the lateral fusiform gyrus. Proceedings of the National Academy of Sciences 117:23011–23020.

Raz N, Amedi A, Zohary E (2005) V1 activation in congenitally blind humans is associated with episodic retrieval. Cereb Cortex 15:1459–1468.

Reddy L, Tsuchiya N, Serre T (2010) Reading the mind’s eye: decoding category information during mental imagery. Neuroimage 50:818–825.

Reich L, Szwed M, Cohen L, Amedi A (2011) A ventral visual stream reading center independent of visual experience. Curr Biol 21:363–368.

Renier L, De Volder AG, Rauschecker JP (2014) Cortical plasticity and preserved function in early blindness. Neurosci Biobehav Rev 41:53–63.

Ricciardi E, Handjaras G, Pietrini P (2014) The blind brain: How (lack of) vision shapes the morphological and functional architecture of the human brain. Experimental Biology and Medicine.

Ricciardi E, Bonino D, Pellegrini S, Pietrini P (2013) Mind the blind brain to understand the sighted one! Is there a supramodal cortical functional architecture? Neuroscience & Biobehavioral Reviews.

Rockland KS, Van Hoesen GW (1994) Direct temporal-occipital feedback connections to striate cortex (V1) in the macaque monkey. Cereb Cortex 4:300–313.

Rockland KS, Ojima H (2003) Multisensory convergence in calcarine visual areas in macaque monkey. Int J Psychophysiol 50:19–26.

Röder B, Rösler F, Neville HJ (2000) Event-related potentials during auditory language processing in congenitally blind and sighted people. Neuropsychologia 38:1482–1502.

Röder B, Stock O, Bien S, Neville H, Rösler F (2002) Speech processing activates visual cortex in congenitally blind humans. Eur J Neurosci 16:930–936.

Rohe T, Noppeney U (2016) Distinct Computational Principles Govern Multisensory Integration in Primary Sensory and Association Cortices. Current Biology 26:509–514.

Roth ZN, Ryoo M, Merriam EP (2020) Task-related activity in human visual cortex. PLOS Biology 18:e3000921.

Sadato N, Pascual-Leone A, Grafman J, Deiber MP, Ibanez V, Hallett M (1998) Neural networks for Braille reading by the blind. Brain 121:1213–1229.

Sadato N, Pascual-Leone A, Grafman J, Ibanez V, Deiber MP, Dold G, Hallett M (1996) Activation of the primary visual cortex by Braille reading in blind subjects. Nature 380:526–528.

Saenz M, Lewis LB, Huth AG, Fine I, Koch C (2008) Visual Motion Area MT+/V5 Responds to Auditory Motion in Human Sight-Recovery Subjects. J Neurosci 28:5141–5148.

Saygin ZM, Osher DE, Koldewyn K, Reynolds G, Gabrieli JD, Saxe RR (2011) Anatomical connectivity patterns predict face selectivity in the fusiform gyrus. Nat Neurosci 15:321–327.

Saygin ZM, Osher DE, Norton ES, Youssoufian DA, Beach SD, Feather J, Gaab N, Gabrieli JDE, Kanwisher N (2016) Connectivity precedes function in the development of the visual word form area. Nat Neurosci advance online publication:1250–1255.

Senden M, Emmerling TC, van Hoof R, Frost MA, Goebel R (2019) Reconstructing imagined letters from early visual cortex reveals tight topographic correspondence between visual mental imagery and perception. Brain Structure and Function.

Sereno MI, Dale AM, Reppas JB, Kwong KK, Belliveau JW, Brady TJ, Rosen BR, Tootell RB (1995) Borders of multiple visual areas in humans revealed by functional magnetic resonance imaging. Science 268:889–893.

Seydell-Greenwald A, Chambers CE, Ferrara K, Newport EL (2020) What you say versus how you say it: Comparing sentence comprehension and emotional prosody processing using fMRI. NeuroImage 209:116509.

Shaywitz BA, Shaywitz SE, Pugh KR, Fulbright RK, Skudlarski P, Mencl WE, Constable RT, Marchione KE, Fletcher JM, Klorman R, Lacadie C, Gore JC (2001) The Functional Neural Architecture of Components of Attention in Language-Processing Tasks. NeuroImage 13:601–612.

Shimony JS, Burton H, Epstein AA, McLaren DG, Sun SW, Snyder AZ (2006) Diffusion tensor imaging reveals white matter reorganization in early blind humans. Cereb Cortex 16:1653–1661.

Shu N, Li J, Li K, Yu C, Jiang T (2009) Abnormal diffusion of cerebral white matter in early blindness. Hum Brain Mapp 30:220–227.

Slotnick SD, Thompson WL, Kosslyn SM (2005) Visual Mental Imagery Induces Retinotopically Organized Activation of Early Visual Areas. Cerebral Cortex 15:1570–1583.

Stănişor L, van der Togt C, Pennartz CMA, Roelfsema PR (2013) A unified selection signal for attention and reward in primary visual cortex. Proceedings of the National Academy of Sciences 110:9136–9141.

Striem-Amit E, Amedi A (2014) Visual Cortex Extrastriate Body-Selective Area Activation in Congenitally Blind People Seeing by Using Sounds. Curr Biol 24:687–692.

Striem-Amit E, Vannuscorps G, Caramazza A (2018a) Plasticity based on compensatory effector use in the association but not primary sensorimotor cortex of people born without hands. Proceedings of the National Academy of Sciences 115:7801–7806.

Striem-Amit E, Cohen L, Dehaene S, Amedi A (2012) Reading with sounds: preserved functional specialization in the ventral visual cortex of the congenitally blind. In: Neuroscience. New Orleans, USA.

Striem-Amit E, Wang X, Bi Y, Caramazza A (2018b) Neural Representation of Visual Concepts in People Born Blind. Nature Communications 9:5250.

Striem-Amit E, Ovadia-Caro S, Caramazza A, Margulies DS, Villringer A, Amedi A (2015) Functional connectivity of visual cortex in the blind follows retinotopic organization principles. Brain 138:1679–1695.

Sturm W, Willmes K (2001) On the functional neuroanatomy of intrinsic and phasic alertness. Neuroimage 14:S76–84.

Takahashi E, Folkerth RD, Galaburda AM, Grant PE (2012) Emerging Cerebral Connectivity in the Human Fetal Brain: An MR Tractography Study. Cerebral Cortex 22:455–464.

van den Hurk J, Van Baelen M, Op de Beeck HP (2017) Development of visual category selectivity in ventral visual cortex does not require visual experience. Proceedings of the National Academy of Sciences 114:E4501–E4510.

Vetter P, Smith FW, Muckli L (2014) Decoding Sound and Imagery Content in Early Visual Cortex. Curr Biol 24:1256–1262.

Vetter P, Bola Ł, Reich L, Bennett M, Muckli L, Amedi A (2020) Decoding Natural Sounds in Early “Visual” Cortex of Congenitally Blind Individuals. Current Biology.

Vigneau M, Beaucousin V, Hervé PY, Duffau H, Crivello F, Houdé O, Mazoyer B, Tzourio-Mazoyer N (2006) Meta-analyzing left hemisphere language areas: phonology, semantics, and sentence processing. Neuroimage 30:1414–1432.

Wandell BA, Winawer J (2011) Imaging retinotopic maps in the human brain. Vision Research 51:718–737.

Wandell BA, Dumoulin SO, Brewer AA (2007a) Visual Field Maps in Human Cortex. Neuron 56:366–383.

Wandell BA, Dumoulin SO, Brewer AA (2007b) Visual field maps in human cortex. Neuron 56:366–383.

Wang D, Qin W, Liu Y, Zhang Y, Jiang T, Yu C (2013) Altered resting-state network connectivity in congenital blind. Human Brain Mapping 35:2573–2581.

Westerhausen R, Moosmann M, Alho K, Belsby SO, Hämäläinen H, Medvedev S, Specht K, Hugdahl K (2010) Identification of attention and cognitive control networks in a parametric auditory fMRI study. Neuropsychologia 48:2075–2081.

Wolmetz M, Poeppel D, Rapp B (2010) What Does the Right Hemisphere Know about Phoneme Categories? Journal of Cognitive Neuroscience 23:552–569.

Yarkoni T, Poldrack RA, Nichols TE, Van Essen DC, Wager TD (2011) Large-scale automated synthesis of human functional neuroimaging data. Nature Methods 8:665–670.

Yu C, Shu N, Li J, Qin W, Jiang T, Li K (2007) Plasticity of the corticospinal tract in early blindness revealed by quantitative analysis of fractional anisotropy based on diffusion tensor tractography. Neuroimage 36:411–417.

Yu C, Liu Y, Li J, Zhou Y, Wang K, Tian L, Qin W, Jiang T, Li K (2008) Altered functional connectivity of primary visual cortex in early blindness. Hum Brain Mapp 29:533–543.

Zekveld AA, Heslenfeld DJ, Festen JM, Schoonhoven R (2006) Top–down and bottom–up processes in speech comprehension. NeuroImage 32:1826–1836.

